# Cortical multi-area model with joint excitatory-inhibitory clusters accounts for spiking statistics, inter-area propagation, and variability dynamics

**DOI:** 10.1101/2024.01.30.577979

**Authors:** Jari Pronold, Aitor Morales-Gregorio, Vahid Rostami, Sacha J. van Albada

## Abstract

The primate brain uses billions of interacting neurons to produce macroscopic dynamics and behavior, but current methods only allow neuroscientists to investigate a subset of the neural activity. Computational modeling offers an alternative testbed for scientific hypotheses, by allowing full control of the system. Here, we test the hypothesis that local cortical circuits are organized into joint clusters of excitatory and inhibitory neurons by investigating the influence of this organizational principle on cortical resting-state spiking activity, inter-area propagation, and variability dynamics. The model represents all vision-related areas in one hemisphere of the macaque cortex with biologically realistic neuron densities and connectivities, expanding on a previous unclustered model of this system. Each area is represented by a square millimeter microcircuit including the full density of neurons and synapses, avoiding downscaling artifacts and testing cortical dynamics at the natural scale. We find that joint excitatory-inhibitory clustering normalizes spiking activity statistics in terms of firing rate distributions and inter-spike interval variability. A comparison with data from cortical areas V1, V4, FEF, 7a, and DP shows that the clustering enables the resting-state activity of especially higher cortical areas to be better captured. In addition, we find that the clustering supports signal propagation across all areas in both feedforward and feedback directions with reasonable latencies. Finally, we also show that localized stimulation of the clustered model quenches the variability of neural activity, in agreement with experimental observations. We conclude that joint clustering of excitatory and inhibitory neurons is a likely organizational principle of local cortical circuits, supporting resting-state spiking activity statistics, inter-area propagation, and variability dynamics.

## Introduction

Most studies in computational neuroscience focus on either the local or global circuitry while neglecting the interactions across scales. Recent studies bridge these scales by combining local and global circuits into multi-scale models enabling cortical simulations at neuronal and synaptic resolution (Schmidt et al., 2018a,b). This approach raises new questions compared to the study of isolated local circuits, for example: How can a hierarchically organized spiking neural network with realistic activity statistics support reliable signal propagation across areas? Reliable signal propagation is considered to be one of the four key properties of a candidate neural code (Perkel and Bullock, 1968; Kumar et al., 2010). Most previous studies have made simplifications, such as considering only strictly feedforward networks, assuming all areas to be identical, and not using biological data to constrain cortical connectivity (Diesmann et al., 1999; Deco and Rolls, 2005; Kumar et al., 2008, 2010). These studies identified a major common issue: modeled signals tend to either die out or amplify. Topographic connectivity has been shown to be crucial for signal propagation in spiking neural networks (Zajzon et al., 2019). A recent study achieved signal propagation in data-driven large-scale models with simplified local connectivity (Joglekar et al., 2018), in both population rate models and connected balanced spiking neural networks, albeit without realistic spiking statistics. Here, we present a large-scale model at single-neuron resolution with clustered connectivity, that is able to transmit signals across the cortical hierarchy while preserving biologically realistic dynamics. To avoid otherwise inevitable downscaling artifacts (van Albada et al., 2015), we include the full biological density of neurons and synapses in each local circuit, yielding a model with about 4 million neurons and 24 billion synapses. Although much is known about cortico-cortical and local connections, the structural connectivity is not fully characterized. Structural relations and statistical regularities can be used to fill the gaps in the anatomical data (Schmidt et al., 2018a; van Albada et al., 2022). The population-level connectivity matrix for the vision-related areas of macaque cortex derived from anatomical data by Schmidt et al. (2018a) was found to produce unrealistic activity when simulated with random connectivity below the population level. The predicted connectivity matrix spans six orders of magnitude and has a relatively high uncertainty. Small changes to the connectivity can increase global stability (Schuecker et al., 2017) by uncovering cortical loops critical to global stability, such as that between areas the frontal eye field and dorsolateral prefrontal cortex, ultimately leading to a stable network (Schmidt et al., 2018b). These neuron-level networks are difficult to control due to their size and complexity. Thus, reliably transmitting signals across areas without destabilizing the network is challenging.

To overcome the limitations of random networks and ensure signal transmission, several studies use clustered networks (Amit and Brunel, 1997; Deco and Rolls, 2005; Litwin-Kumar and Doiron, 2012; Mazzucato et al., 2015; Rost et al., 2018; Rostami et al., 2022). Clustered networks involve some form of strengthened connections within clusters and weakened connections across clusters. Deco and Rolls (2005) studied attention using a network of spiking neurons spanning two areas and featuring different clusters of excitatory neurons within the areas. Clustered networks tend to have multiple attractors. Thus, models with clusters of excitatory neurons have been used to explain decision-related activity (Amit and Brunel, 1997; Litwin-Kumar and Doiron, 2012; Mazzucato et al., 2015). Jointly clustering excitatory and inhibitory populations can be used to robustly build winnerless competition into balanced random networks while reproducing biological firing rates, spiking irregularity, and trial-to-trial spike count variability from in vivo recordings (Rost et al., 2018; Rostami et al., 2022). Furthermore, excitatory-inhibitory (EI) clustering can generate robust multistability with local balance for a wider range of network sizes and parameters than purely excitatory (E) clustering (Schaub et al., 2015; Rost et al., 2018; Najafi et al., 2020; Rostami et al., 2022).

Ample evidence exists that synaptic connections between excitatory neurons are clustered and not uniform (Song et al., 2005; Perin et al., 2011). For example, Song et al. (2005) found that, in the visual system, bidirectional and clustered three-neuron connection motifs occur significantly more often than in a random graph based on a pairwise connection probability alone. Furthermore, such clusters receive similar visual feedforward input and thus could form fine-scale functional groups (Yoshimura et al., 2005; Ko et al., 2011). Clusters can be identified in the neocortex as ensembles of highly active, interconnected cells and might encode sensory information by high firing rates (Yassin et al., 2010). More recent anatomical and physiological findings suggest that inhibitory neurons and their connectivity also have a high degree of specificity (Xue et al., 2014; Lee et al., 2014; Morishima et al., 2017; Arkhipov et al., 2018; Khan et al., 2018; Znamenskiy et al., 2018; Shin et al., 2019; Najafi et al., 2020). For instance, in reciprocally connected pairs of inhibitory and excitatory neurons in the mouse visual cortex there is a positive correlation between the strength of the excitatory and the inhibitory synapses (Znamenskiy et al., 2024). These studies suggest that the networks can form strong local interconnected clusters consisting of both excitatory and inhibitory cells. All in all, clustered connectivity is a common feature of the brain that can be computationally advantageous, but its effects on large-scale dynamics remain to be elucidated.

In this work, we introduce a clustered connectivity scheme (Fig. 1) in a biologically constrained model of macaque cortex (Schmidt et al., 2018a,b). We validate our clustered model by comparing its simulated activity with the previous unclustered version of the model, as well as with resting-state spiking activity across several cortical areas (V1, V4, FEF, 7a, and DP). We find that the clustered model supports plausible activity statistics in terms of firing rate distributions and inter-spike interval variability. We show that the clustered connectivity scheme improves upon the original model particularly in terms of the activity statistics of the higher cortical areas FEF, 7a, and DP. Most importantly, we find that the clustered model can transmit signals across all areas in both feedforward and feedback directions. Upon stimulation of V1, the activity propagates through the entire model with plausible response latencies. Finally, we also show that the stimulation of the model quenches the variability of neural activity, in agreement with experimental observations. All in all, we show how joint clustering of excitatory and inhibitory neurons can support plausible resting-state spiking activity statistics, inter-area signal propagation, and variability dynamics upon stimulation in a multi-area cortical network. The model can function as a testbed for further studies of cortical spiking dynamics requiring inter-areal signal propagation.

**Figure 1:**
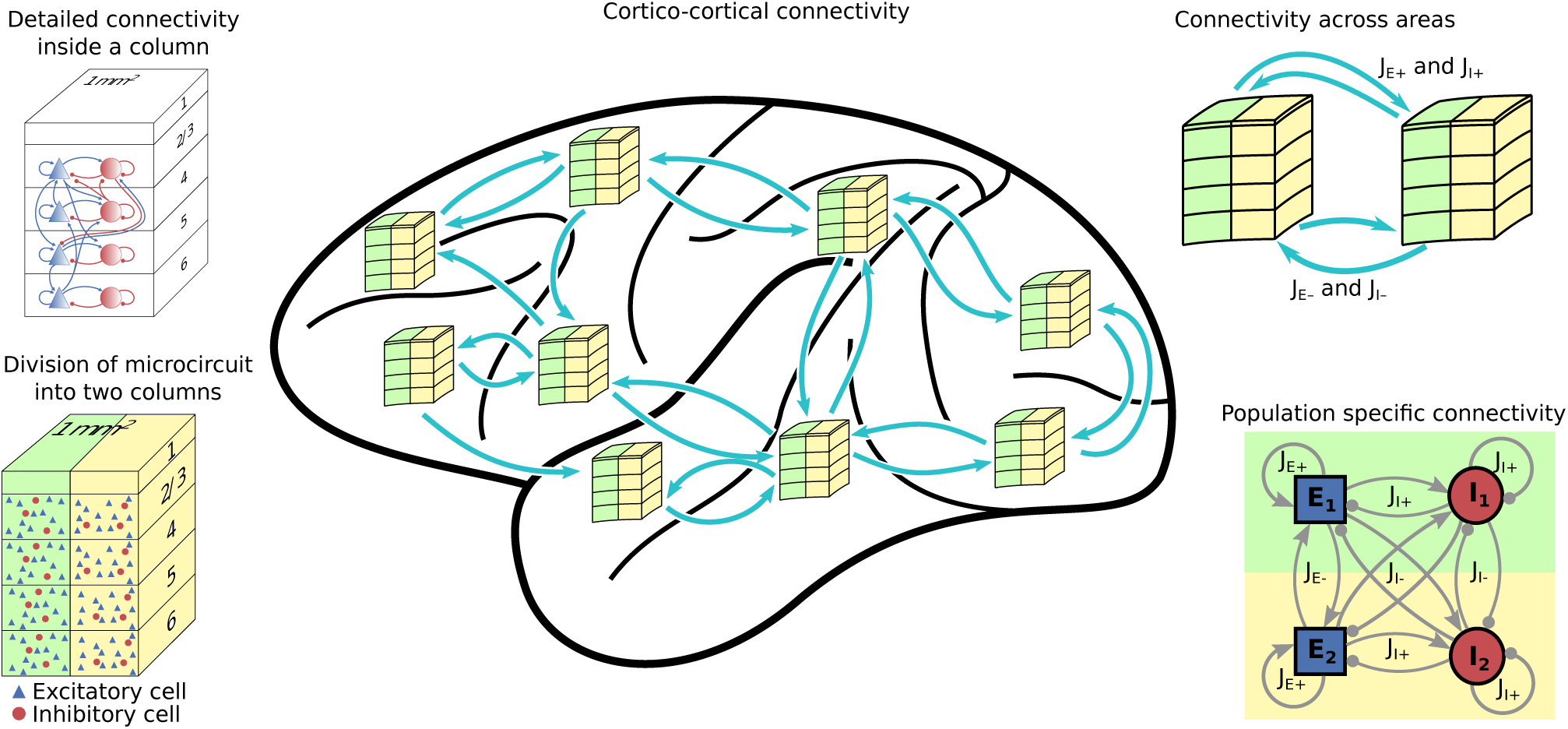
Overview of the model. The basic building block of the areas of the model is the microcircuit and its intricate connectivity (Potjans and Diesmann, 2014). All areas are split into *Q* clusters of equal size (here, *Q* = 2 is shown). Every column in every area has a counterpart in every other area. The synaptic weights are scaled inside and across columns, both locally and across areas. The scaling depends on whether the same or different columns connect with each other. If the connection is established within the same (or, for inter-area connections, corresponding) cluster, the weights are strengthened with the factors *J_E_*_+_ and *J_I_*_+_. Otherwise the connections are weakened by the factors *J_E−_* and *J_I−_*. This is illustrated on the bottom right with a two-population network for simplicity.

## Results

### A clustered multi-area model of the macaque cortex

To study the effect of network clustering in macaque cortex we use a previously developed multi-area model of the vision-related areas (Schuecker et al., 2017; Schmidt et al., 2018a,b), see Methods for a detailed account of the model construction. In short, the multi-area model is a multi-scale spiking network model of all vision-related areas in one hemisphere of macaque cortex with neuronal and synaptic resolution. It integrates experimental data on cortical architecture and connectivity into a comprehensive network and relates cortical connectivity to its dynamics. The local circuitry features four cortical layers and uses the microcircuit of Potjans and Diesmann (2014) as a blueprint. The cortico-cortical connectivity is based on axonal tracing data (Bakker et al., 2012), including quantitative and layer-specific retrograde tracing data (Markov et al., 2014b,a). We extend this model by subdividing every area into *Q* clusters. Both inside and across areas, synapses within clusters are strengthened, and synapses between different clusters are weakened. Fig. 1 schematically shows the network construction for *Q* = 2.

### Unrealistic aspects of the dynamics of the original multi-area model

As a baseline for comparison, we first simulated the spiking activity of the original unclustered model (Schmidt et al., 2018a,b). Fig. 2A shows raster plots for areas V1, V2, V4, 7a, DP, and FEF when the model is in the metastable state (see Methods). V1, V2, and FEF show largely asynchronous irregular activity, while the vertical stripes for V4, 7a, and DP indicate high synchrony. Neurons in areas 7a and DP fire at a high rate, especially in layers 4 and 5 of area 7a.

**Figure 2:**
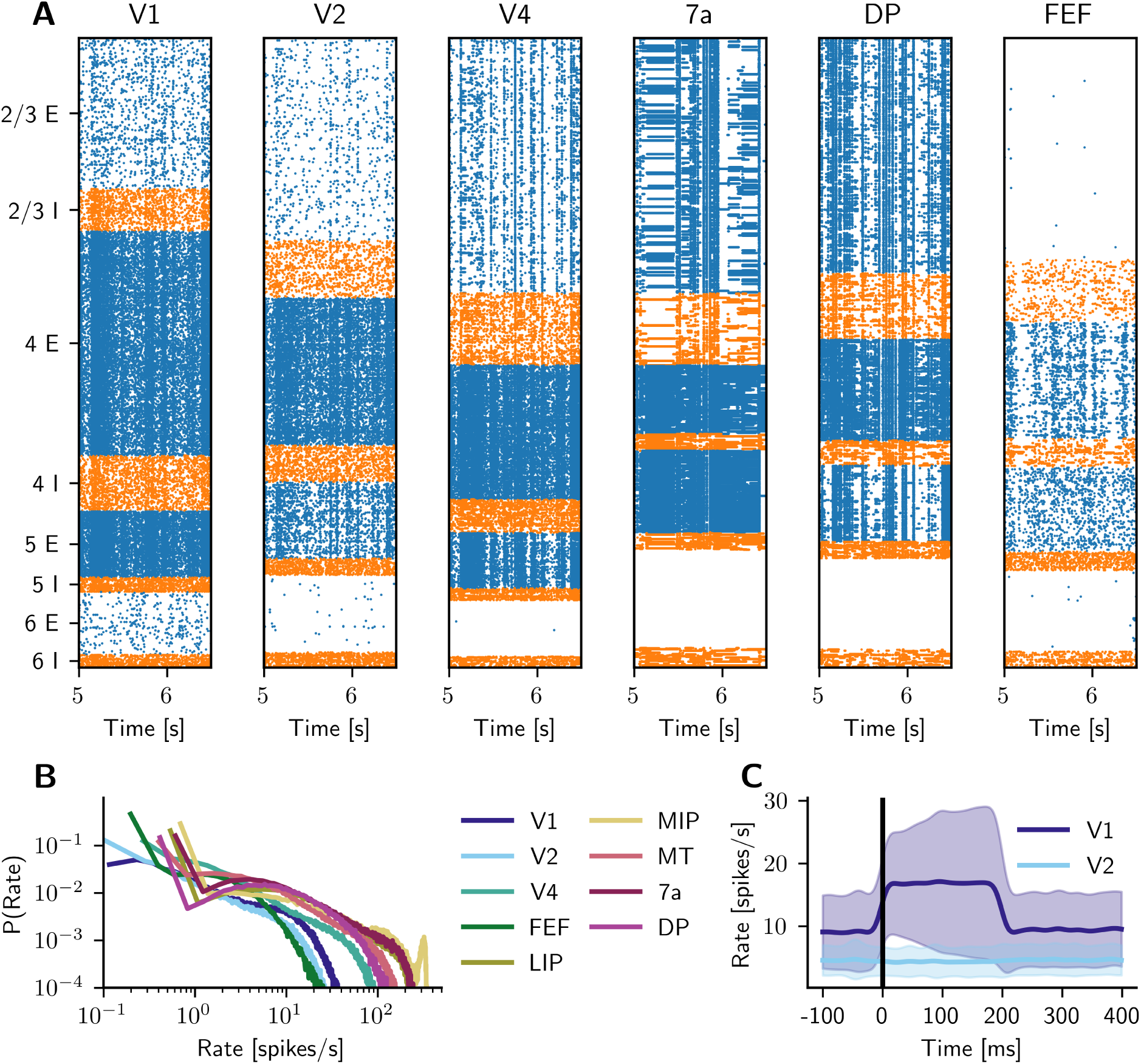
Dynamics of the original model. (**A**) Raster plot of spiking activity of 2% of the neurons in areas V1, V2, V4, 7a, DP, and FEF during resting state. Blue: excitatory neurons, orange: inhibitory neurons. **(B)** Distribution of spike rates across all layers and populations for several cortical areas. **(C)** Propagation of a stimulation of 200 ms of 30 spikes*/*s from area V1 to V2. The firing rates are averaged over 100 trials and convolved with a Gaussian kernel with a width of 10 ms. Standard deviation (±) at each time point is indicated by the shaded regions.

Fig. 2B shows the distribution of firing rates for selected areas. Area MIP shows an unrealistic firing peak at around 300 spikes*/*s. Areas DP, MT, LIP, and 7a show a high density of firing rates well above 100 spikes*/*s. The other areas fire at more reasonable rates. The raster plots of 7a and DP in Fig. 2A suggest that population 5E spikes excessively while population 6E stays completely silent.

To assess targeted signal propagation between areas, we stimulate V1 and observe the resulting firing rates in V2 in Fig. 2C. The stimulation is a brief pulse of 200 ms with a rate of 30 spikes*/*s. We recorded the instantaneous firing rate from V1 and V2 with a bin size of 1 ms and convolved it with a Gaussian kernel of 10 ms width. The stimulation was repeated every second, providing 100 stimulation trials. Fig. 2C shows the mean (solid line) and standard deviation (shading) across trials. No detectable signal arrives in area V2 in response to the V1 stimulus, even though most of the cortico-cortical connections to V2 originate from V1. The large standard deviation in the firing rates of area V1 indicates that the same stimulus can have vastly different effects on the dynamics of V1. As shown in Schmidt et al. 2018b, signals do propagate through the network spontaneously, but it appears difficult to control their directions and strengths.

### Clustering supports realistic spiking dynamics and signal propagation

To address the lack of signal propagation in the original multi-area model, we introduce a connectivity structure that clusters areas into columns and alters the synaptic weights. The altered synaptic weights emphasize connections between the same columns and weaken all other connections (see Methods Clustered multi-area model of macaque visual cortex). We simulated a clustered model with *Q* = 50 clusters. Fig. 3A shows raster plots for V1, V2, V4, 7a, DP, and FEF in the condition without transient stimulation, mimicking the resting state. The overall firing behavior has considerably changed with respect to the original unclustered model: In every area, some clusters are more active than others, displaying activity akin to up states. Especially in FEF, there is frequent switching between active clusters. Furthermore, the vertical stripes are gone, and spiking activity in areas 7a and DP is much more plausible. Fig. 3B shows the distribution of firing rates for selected cortical areas. All areas have a comparable firing rate distribution, and no area spikes excessively, in agreement with experimentally measured activity levels (Shinomoto et al., 2003, 2009; Morales-Gregorio et al., 2020).

**Figure 3:**
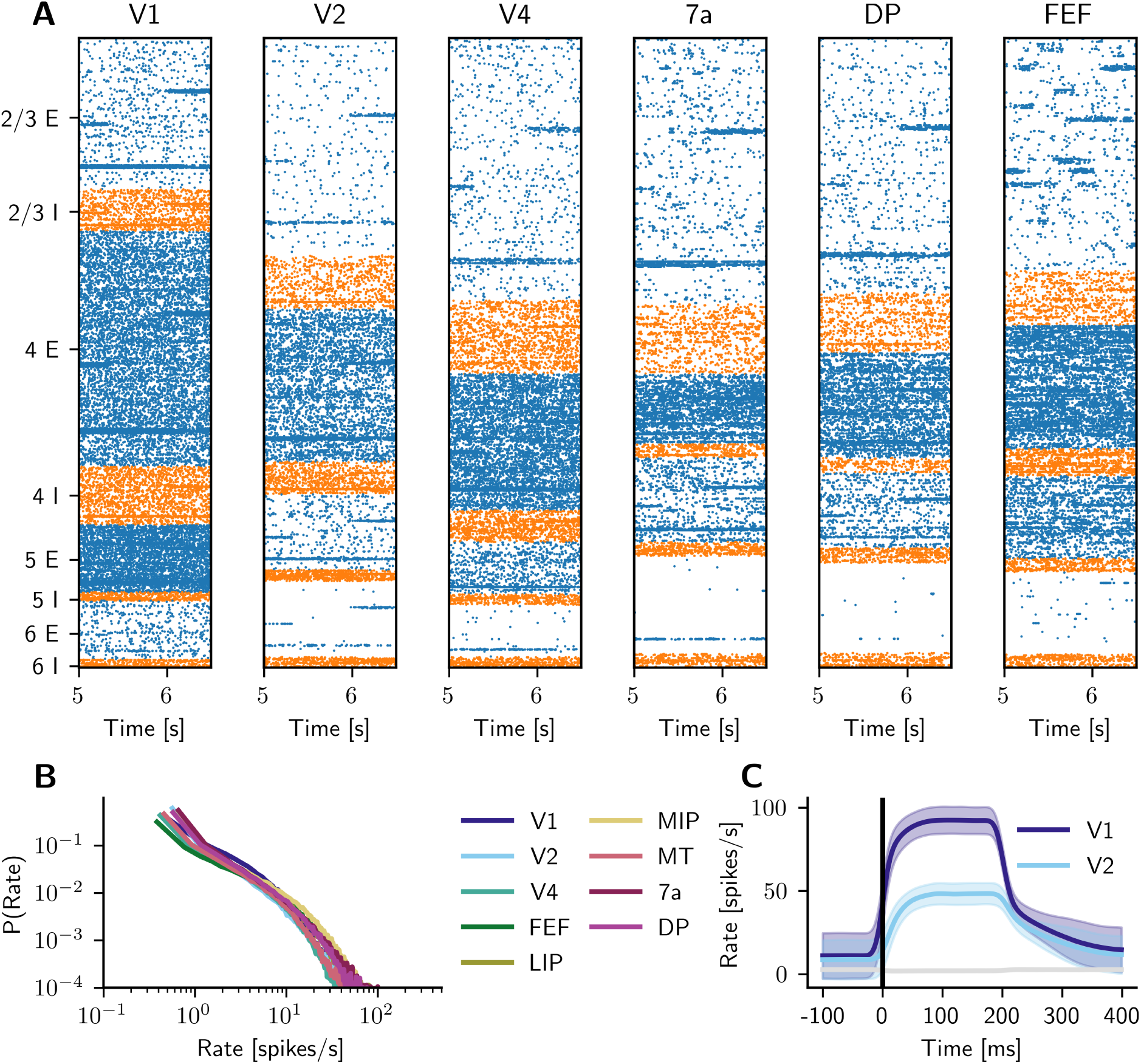
Dynamics of the clustered model. (**A**) Raster plot of spiking activity of 2% of the neurons in areas V1, V2, V4, 7a, DP, and FEF during the simulated resting state. Neurons are ordered according to their cluster membership. Blue: excitatory neurons, orange: inhibitory neurons. **(B)** Distribution of spike rates across all layers and populations for several cortical areas. **(C)** Propagation of a stimulation of 200 ms of 30 spikes*/*s from the stimulated cluster in area V1 to the corresponding cluster in V2. The firing rates are averaged over 100 trials and convolved with a Gaussian kernel with a width of 10 ms. Standard deviation (±) at each time point is indicated by the shaded regions. The gray line shows the firing rate of all non-stimulated clusters in area V1.

In order to study signal propagation, we provided a stimulation of 200 ms with a rate of 30 spikes*/*s to a given cluster in V1, repeated every second. Fig. 3C shows the propagation of the signal from V1 to the corresponding cluster in V2. A clearly detectable signal arrives in V2 briefly after V1 stimulation. The standard deviation of firing rates in areas V1 and V2 is smaller than in the original model (see Fig. 2C), indicating that the same stimulus has a more predictable effect on the firing rates. The gray line in Fig. 3C shows the firing rate of all other clusters in V1. Thus, the firing rates in the non-stimulated clusters remain low and are slightly suppressed during stimulation.

We have thus shown that the clustered model has more realistic spiking activity with a consistent firing rate distribution across areas and can reliably propagate a stimulus from V1 to V2.

### Comparison of single-neuron spiking statistics with experimental recordings

We compare the simulated data from the original model and the clustered model with new spiking neuron data (see methods Experimental data). The experimental data consist of recordings from V1 layers 5/6, V4 layers 2/3 and 5/6, FEF layers 2/3, 7a layers 5/6 and DP layers 5/6. Fig. 4 compares the distributions of the coefficient of variation of the inter-spike intervals (CV ISI), the revised local variation (LvR; Shinomoto et al., 2009), and firing rate in all areas. In 7a and DP, the two experimental lines correspond to different recording sessions. To match the number of neurons from the experimental data, we randomly sample the same number of spike trains from the simulated data (*N* = 100 realizations) and plot the mean (lines) and standard deviation (shadings). The clustered model (*Q* = 50) matches the experimental data better than the original model for all data sets except V4 L2/3. It especially outperforms the original model for V4 L5/6, 7a, and DP. The CV ISI distribution in 7a of the original model (shown as an inset) is centered around unrealistically high values. In DP, the original CV ISI distribution is also shifted to the right, but not as strongly. The distributions of all three measures, CV ISI, LvR, and firing rate, match the experimental data better in the clustered than in the original model. In the lower panel, we show the Kolmogorov-Smirnov distance between the experimental and simulated distributions for the CV ISI, LvR, and firing rate for different numbers of clusters *Q*. In the case of the CV ISI distribution, V1, 7a, and DP profit from clustering, while V4 and FEF initially worsen a little bit but recover at *Q* = 50. In the case of the CV ISI distribution, V1, 7a, and DP profit from clustering, while V4 and FEF initially worsen slightly but recover at *Q* = 50. For the LvR distribution, most areas appear unaffected by clustering, while 7a and DP profit from clustering, and V4 worsens. The agreement of the rate distribution does not change much with *Q*; only the V4 agreement declines.

**Figure 4:**
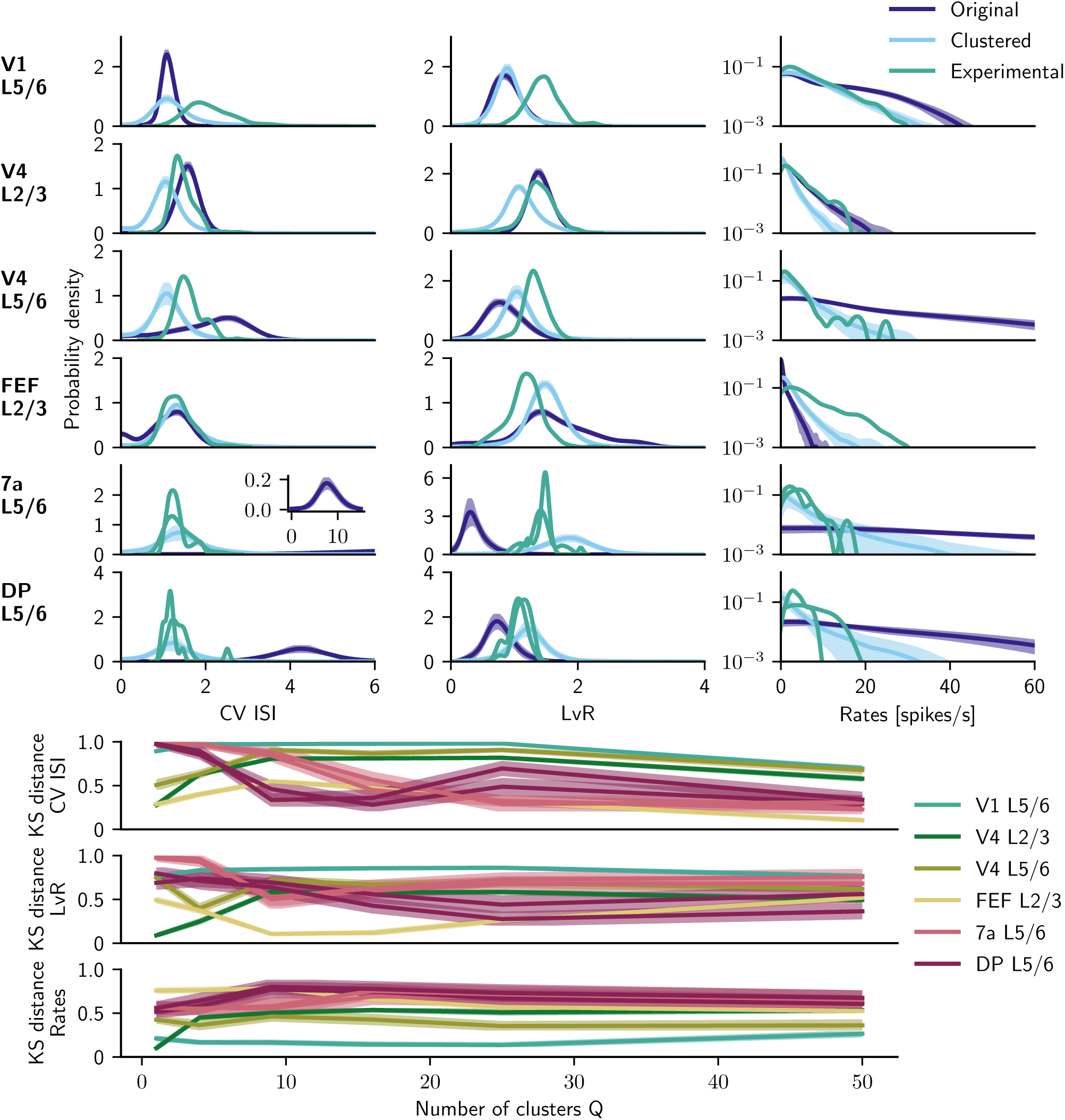
Comparison of single-neuron spiking statistics from clustered model with experimental recordings. Upper panel: Probability densities of CV ISI, LvR, and rate distributions for area V1 layers 5 and 6, area V4 layers 2 and 3, area V4 layers 5 and 6, area FEF layers 2 and 3, 7a layers 5 and 6, and DP layers 5 and 6. The experimental data in 7a and DP contain two measurements from two independent recording sessions. The clustered model uses *Q* = 50 clusters. Inset in 7a, layer 5/6 shows the CV ISI of the original model. Lower panel: Kolmogorov-Smirnov distance of the CV ISI, LvR, and rate distribution comparing experimental and simulated data for different numbers of clusters *Q*. The left data point at *Q* = 1 equals the original, metastable model of Schmidt et al. (2018b).

### Network-wide signal propagation in feedforward and feedback directions

We simulate the response of the clustered model to a pulse of 200 ms and 30 spikes*/*s applied either to one cluster in primary visual cortex (area V1) or to one cluster in the frontal eye field (FEF). Fig. 5 shows the firing rates in the stimulated cluster for feedforward and feedback propagation. Areas are sorted according to their distance to the stimulated area, measured as the shortest possible path between the areas without crossing the cortical surface (Bojak et al., 2011). The coloring of the area names corresponds to different modules of the network determined using the map equation method (Rosvall et al., 2009) applied to the structural connectivity as described in Schmidt et al. (2018b). The firing rates are averaged over 100 trials and convolved with a Gaussian kernel with a width of 10 ms.

**Figure 5:**
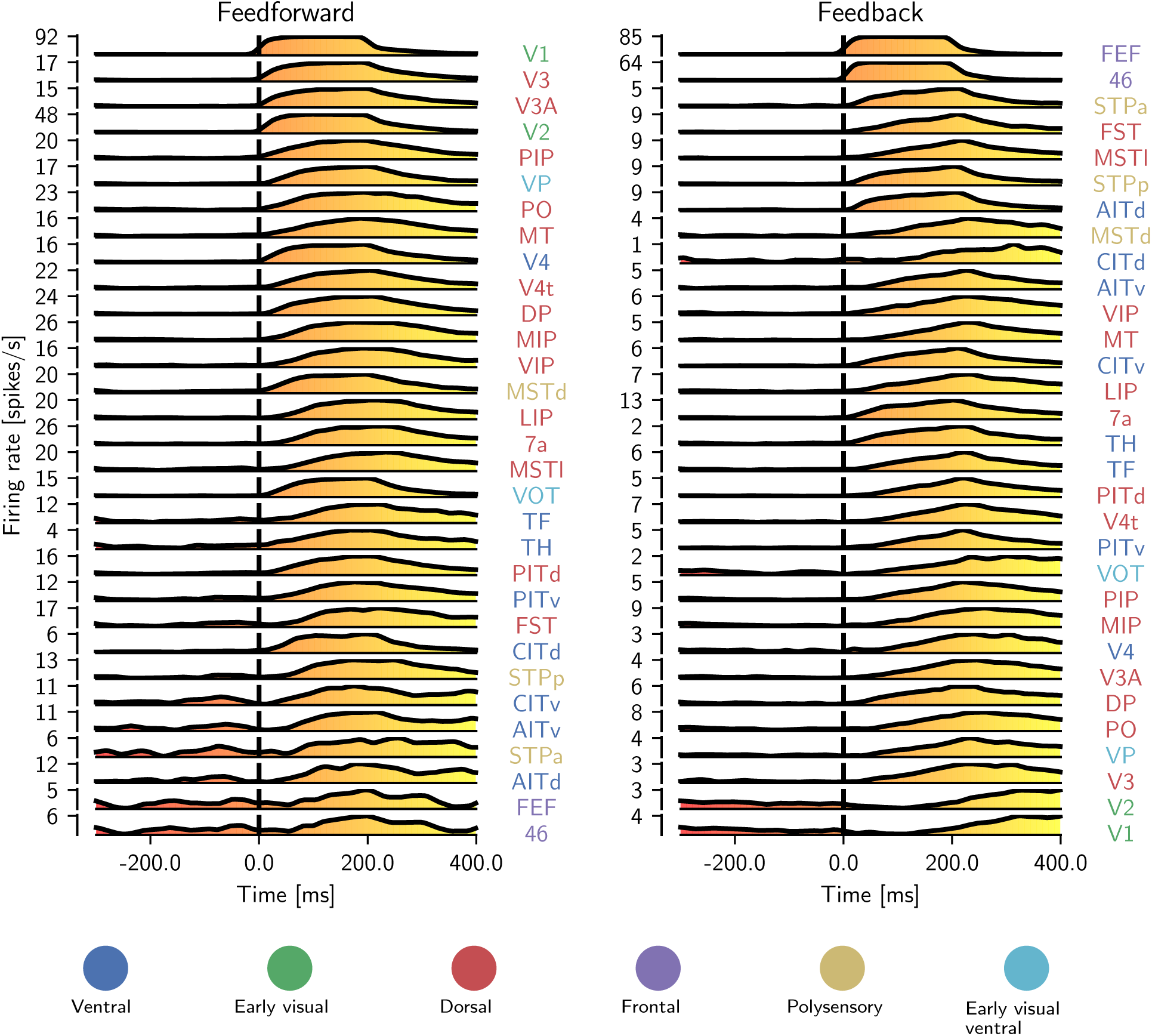
Clustered connectivity enables signal propagation across the entire network, in both the feedforward and feedback directions. A stimulation of 200 ms and 30 spikes*/*s was applied to one out of 50 clusters at time point 0 in area V1 in the feedforward case and in area FEF in the feedback case. The simulations consisted of 100 trials of 1 s duration each. The firing rates are convolved with a Gaussian kernel with a width of 10 ms. Areas are sorted according to shortest path lengths with respect to the stimulated area—either V1 or FEF. The color gradient under the curve represents time. Coloring of the area name labels corresponds to modules of the area-level network identified using the map equation method (Rosvall et al., 2009) as described in Schmidt et al. (2018b). Area MDP is not shown, as it does not have any incoming connections from the rest of the modeled network.

The response to V1 stimulation propagates through the network in the feedforward direction from lower to higher areas in the visual hierarchy. Early visual areas (i.e., those close to the sensory periphery) and dorsal stream areas become active first. Ventral stream and polysensory areas follow these, and the frontal areas are the last to respond. The activation timing coincides well with the sorting according to the shortest paths.

In contrast, the response to FEF stimulation propagates in the feedback direction. In this experiment, frontal areas become active first, followed by polysensory, ventral stream, and dorsal stream areas. Finally, early visual areas respond. We can also observe a relation between distance to the stimulated area and response time, although weaker than in the feedforward case.

In both cases, farther areas respond later to the stimulus and have, in general, a weaker time-integrated response. All areas in the model show a response to both the feedforward and feedback stimulus, with the notable exception of MDP, which has no incoming connections in the model (from the areas included in it) and is therefore not shown in Fig. 5. In summary, the clustered connectivity enables signal propagation throughout the cortex.

### Response latencies in the clustered model

To quantify the speed and effectiveness of the signal propagation, we measured the response latencies in the clustered model. We first used the Poisson surprise algorithm to detect which neuronal responses occurred due to the stimulus and not just by random chance (see methods Response latency measurement via Poisson surprise). In Fig. 6A–D, we show the instantaneous firing rate of all nonrandom neuron responses in four areas (V1, V2, V4, and FEF) and the spike train raster plot of a single sample neuron across trials. Both the firing rate and the raster plot show high activity during the stimulation that decays after the stimulus is removed. The firing rate shows that neurons in V1 respond quickly to the stimulus, whereas in the other areas, the maximum firing rate is reached after a small delay, as also shown in Fig. 5. For all neurons, we measured the time to the first nonrandom spike—the time between stimulus onset and the first nonrandom spike in the corresponding cluster—and depict its cumulative distribution in Fig. 6E. Fig. 6F shows the response latencies, defined as the time point when half of the neurons have become active, compared against experimental data (Schmolesky et al., 1998; Barash et al., 1991; Bushnell et al., 1981; Chafee and Goldman-Rakic, 1998; Robinson et al., 1978; Lamme and Roelfsema, 2000). In the experimental data, the timings are reported from the moment a stimulation was presented to the monkey, whereas in the simulation, we directly stimulate V1. Thus, to have the same frame of reference, we subtracted the experimentally measured latency of V1 from the experimental response latencies. Areas V2, V4, and TF have similar response latencies in our model and in the experimental data. Also, model areas AITd and AITv, both of which overlap with area TEa for which the latency was measured experimentally, on average have a latency comparable to TEa. The remaining areas have a longer response latency in our model than in the *in vivo* experiments.

**Figure 6:**
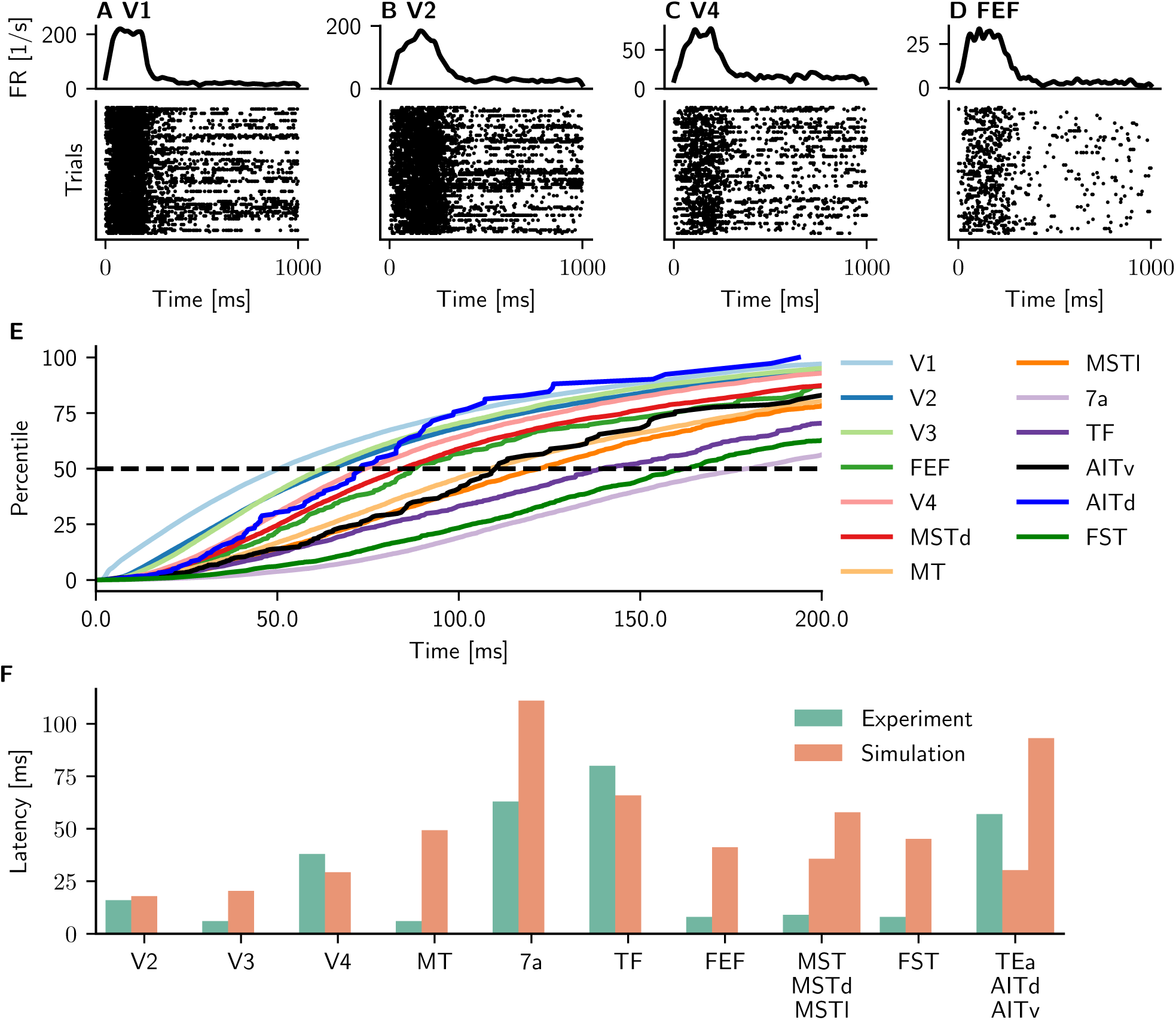
Response latencies in the clustered model. (**A–D**) Neuronal responses of single neurons after stimulation of V1. The shown responses are detected via the Poisson surprise method. Upper panels: Instantaneous firing rate (FR) of all detected neurons for areas V1, V2, V4, and FEF (convolved with a Gaussian kernel with a width of 10 ms). Lower panels: sample raster plot of a single neuron that exhibits a nonrandom response to the stimulus. **(E)** Cumulative distributions of the time to the first nonrandom spike for several cortical areas. The 50th percentile—the time at which half of all neurons within the corresponding cluster have responded to the stimulus—is considered the response latency for that area. **(F)** Comparison of response latencies in experiments (Schmolesky et al., 1998; Barash et al., 1991; Bushnell et al., 1981; Chafee and Goldman-Rakic, 1998; Robinson et al., 1978; Lamme and Roelfsema, 2000) and simulations. The latencies of modeled areas MSTd and MSTl are compared with measurements from area MST; those of modeled areas AITd and AITv are compared with measurements from TEa.

### Quenched neural variability after stimulation

The variability of neuronal activity across trials, measured as the Fano factor, has been shown to decrease after stimulus presentation (Churchland et al., 2010; Rostami et al., 2022). We use the original and clustered models to study whether stimulation reproduces the decrease in neuronal variability. Similarly to the previous simulations, a pulse of 30 spikes*/*s but now lasting 400 ms was applied 50 times every 2 seconds to one cluster in area V1, and we measured the resulting Fano factor (see Methods Neural variability quantification across trials). Fig. 7 depicts the mean-matched Fano factor (solid line) for areas V1, V4, LIP, and MT, along with the standard deviation (shading). In the experimental data, the variability decays as soon as an input is presented, followed by a slow increase and finally stabilizing at a slightly higher value towards the end of the stimulation period. In the original model, the Fano factor of V1 rises sharply when the stimulation is applied. All other areas appear unaffected by the stimulation, further demonstrating the lack of signal propagation. The Fano factor for these areas in the original model is several times larger than the experimental observations, likely due to the strong fluctuations in the simulated activity leading to a high variance across trials. In the clustered model, the stimulation decreases the Fano factor for all areas, albeit not as sharply as in the experimental observations. The Fano factor drops during the stimulation, and in V1 and V4, it reaches a low-value plateau, in contrast to the experimental data, where the Fano factor increases shortly after reaching the minimum.

**Figure 7:**
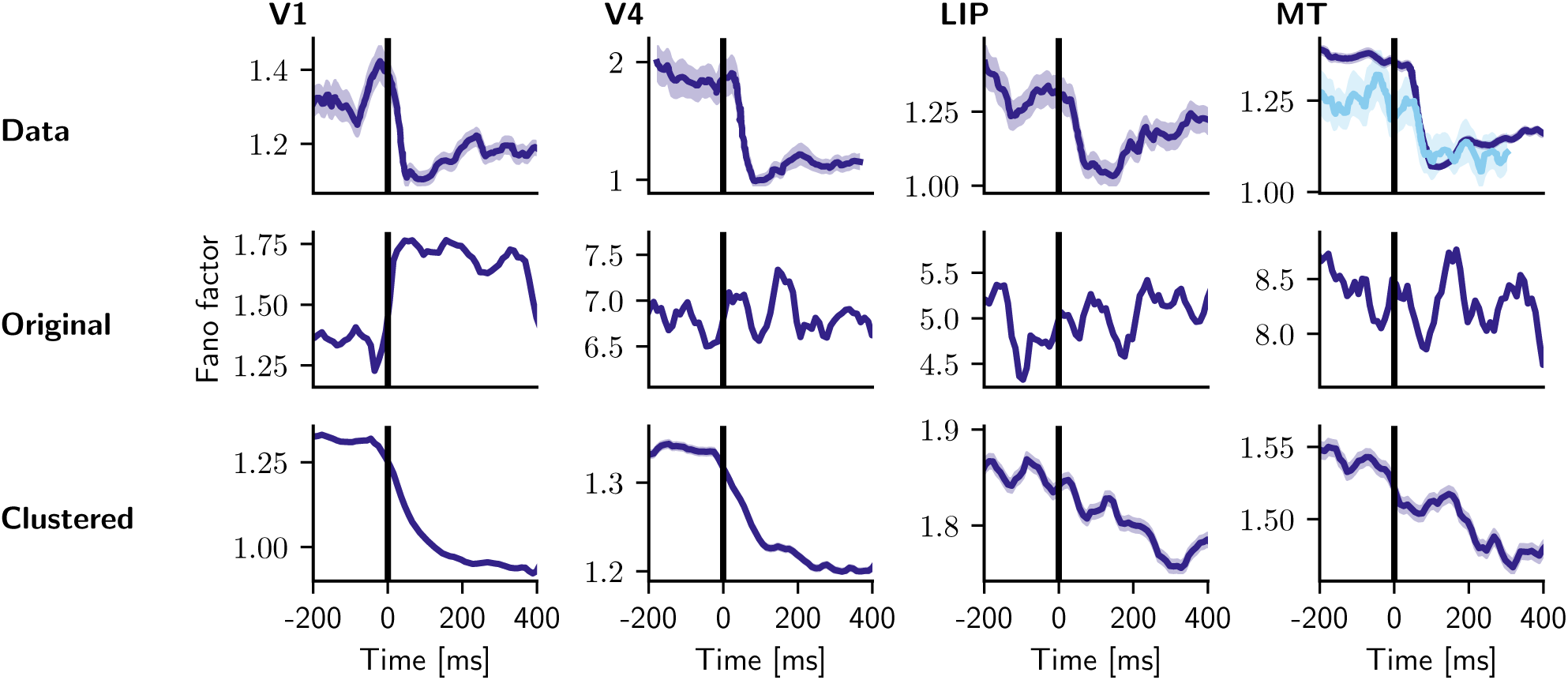
Neural variability changes after stimulation. The evolution of the Fano factor for areas V1, V4, LIP, and MT in experimental data (Churchland et al., 2010), the original model, and the clustered model. MT shows two experimental datasets. Simulated data: a pulse of 400 ms and 30 spikes*/*s was applied to one cluster in primary visual cortex (area V1). In all three panels, Fano factor (FF) is computed in a 50 ms sliding window sliding in steps of 10 ms. Shaded areas show the standard error of the mean.

## Discussion

On the example of a multi-scale model of one hemisphere of macaque visual cortex, we have shown that joint clustering of excitatory and inhibitory neurons helps account for various aspects of cortical dynamics. First, the statistics of ongoing spiking activity in several cortical areas are more realistic compared to an unclustered version of the model (Schmidt et al., 2018a,b). Second, the clustering enables signals to reliably propagate across areas, with response times upon V1 stimulation matching experimental data in several areas. Third, the clustered model reproduces reductions in trial-to-trial variability upon stimulus presentation.

In the original, unclustered model of Schmidt et al. (2018a,b), the simulated V1 spiking activity was compared with parallel spike train recordings from all layers of V1 in lightly anesthetized macaque (Chu et al., 2014b,a). For this area, our model with 50 clusters reproduces the distribution of the spiking irregularity quantified by the coefficient of variation of the inter-spike intervals (CV ISI) somewhat better than the original model (Fig. S1). However, the clustered model has a flatter power spectrum and lower power than both the experimental data and the original model, lacking the population bursts seen in the original model and having somewhat lower firing rates. A partial explanation for this may be that the clustered model is more representative of an awake state rather than an anesthetized state (Hayashi et al., 2014), since the anesthetic ketamine has been found to increase at least low-frequency and gamma power; although it was also found to decrease beta power (Akeju et al., 2016; Schroeder et al., 2016).

In the present study, we compared the spiking activity with data from V1, visual area V4, the frontal eye field (FEF), parietal area 7a, and dorsal prelunate cortex (area DP). Overall, the clustered model fits the experimental data better than the unclustered version, in terms of firing rate distributions and distributions of spiking irregularity quantified by CV ISI and revised local variation (LvR). The main exception is V4 layers 2 and 3, for which the original model performs better. Increasing the number of clusters beyond a few dozen consistently improves the goodness of fit to the experimental data.

The ability to reliably and quickly transmit signals through the brain is critical for implementing a large number of functions. Studying the neuronal basis of complex interactions of feedforward and feedback signals requires a biologically realistic model supporting signal propagation. Joglekar et al. (2018) studied signal transmission in a large-scale model of macaque cortex consisting of population rate models and in a spiking network model. The authors increased cortico-cortical excitatory-to-excitatory and local inhibitory-to-excitatory weights, a scheme they called global balanced amplification following Murphy and Miller (2009). While V1 activation led to signal propagation to some areas in the asynchronous regime, reliable signal propagation across the entire network was only achieved in a synchronized regime. The laminar structure of cortex was neglected and a constant rather than an area– and population-specific connection probability was used. Furthermore, the network was heavily downscaled, containing only 2000 neurons per area, so that an area in their model can be thought of as corresponding roughly to a single cluster in our model. The present study shows how joint clustering of excitatory and inhibitory neurons can support signal propagation across the cortical network at a realistic density of neurons and synapses while maintaining overall asynchronous irregular firing.

The presented clustered model can transmit signals in bottom-up and top-down directions. The signal transmission enabled us to study the response latencies of feedforward signals and compare them with experimental data. In V2 and V4, areas close to V1, the origin of the stimulus, response latencies match well with the reported timings. Also the response latency of area TF and the average latencies of areas AITd and AITv are similar to the experimental findings. To the remaining areas tested, the propagation delay is longer than reported in the literature. We hypothesize that including a pulvinar module could speed up the corresponding signal propagation. The pulvinar connects with most areas of visual cortex (Shipp, 2003; Jones, 2012; Noudoost et al., 2010). Thus, it could decrease latencies by acting as a shortcut linking distant hierarchical levels (Cortes and Van Vreeswijk, 2012; Zajzon and Morales-Gregorio, 2019).

Moreover, including a pulvinar module could facilitate the study of attentional processing: Pulvinar neurons show firing rate modulation with attention (Petersen et al., 1987), and lesions to the pulvinar result in hemispatial neglect toward the contralesional visual field (Petersen et al., 1987; Karnath et al., 2002; Wilke et al., 2010, 2013) and problems in distractor filtering (Desimone et al., 1990), which has also been shown in a computational study (Jaramillo et al., 2019). Attention can be directed by either physical salience of a stimulus (bottom-up) or internal behavioral goals (top-down) (Noudoost et al., 2010). Thus, an extension of the presented model could be used to study the interplay of these two attentional streams and to reproduce experimental findings requiring top-down and bottom-up interactions.

Finally, we studied trial-to-trial variability. An experimental study (Churchland et al., 2010) reports quenching of the Fano factor as a direct result of stimulation. No decline in the Fano factor can be found in the original model because it cannot effectively transmit signals across areas. Moreover, neurons in some areas show an exceptionally high Fano factor. The clustered model, however, displays a clear decline in the Fano factor following the stimulation. The decline lasts as long as the stimulation is active. In contrast, a slowly rising Fano factor follows an initial, low plateau in the experimental data. We hypothesize that using an adaptive neuron model would allow the model to adjust to the stimulation input, and thus, the Fano factor would rise slowly after reaching a minimum.

In a next step, the size of the clusters could be made area-specific. The Kolmogorov-Smirnov distance of the CV ISI (Fig. 4) decreases with the number of clusters. With the current setup, further increasing the number of clusters is not feasible, as the model is already highly computationally intensive. The network construction time jumps from one minute in the original model to seven hours in the clustered model. The increased construction time is due to the increased numbers of nest.Connect calls: The original model has 254 populations connected using 9,116 connect calls. The clustered model consisting of 50 clusters has 50 *·* 254 = 12,700 populations connected by 9,116 *·* 50^2^ = 22,790,000 connect calls. This number could potentially be reduced by implementing a specialized connection routine in the NEST simulator to handle clustered connectivity.

To summarize, we introduce joint excitatory-inhibitory clustering in a biologically based multi-scale spiking model of one hemisphere of macaque vision-related cortex. This connectivity scheme supports inter-area signal propagation and a reduction in trial-to-trial variability upon stimulation, and enabled us to study response latencies. Furthermore, the clustered model reproduces spiking activity statistics in several cortical areas and retains most of the explanatory power of the original model. The clustered model can be used in future studies to elucidate information processing involving bottom-up and top-down interactions and to study the impact of subcortical structures on signal propagation.

## Methods

### Multi-area model of macaque visual cortex

The multi-area model is a multi-scale spiking network model of the vision-related areas in one hemisphere of macaque cortex and relates cortical connectivity to its resting-state dynamics. It integrates cortical architecture and connectivity data into a comprehensive network of 32 areas. Each area consists of the four layers 2/3, 4, 5, and 6, containing one excitatory and one inhibitory population each, and is represented by a patch of 1 mm^2^. Only agranular area TH consists of three layers, as it lacks layer 4. Table 2 summarizes the original and modified model, and Table 3 gives the neuron and synapse parameters. The inter-area (cortico-cortical) connectivity is based on axonal tracing data from the CoCoMac database (Bakker et al., 2012) combined with data from quantitative and layer-specific retrograde tracing experiments (Markov et al., 2014b,a). Local connectivity is based on the microcircuit model of Potjans and Diesmann (2014). The local microcircuit is customized for every area according to the neuronal densities and laminar thicknesses (Schmidt et al., 2018a). Combining local and cortico-cortical connectivity results in a connectivity matrix which is then stabilized using mean-field theory. The stabilization is necessary to arrive at a dynamical state that yields non-vanishing, non-saturating spike rates (Schuecker et al., 2017). The refined connectivity is used to simulate macaque vision-related cortex. By controlling the strength of the cortico-cortical interactions, the model can be poised in a metastable state where simulations reproduce local and cortico-cortical experimental findings (Schmidt et al., 2018b): the V1 single-cell spiking statistics, expressed as firing rates and power spectra, are close to those from recordings in macaque V1. The resulting inter-area functional connectivity patterns match macaque fMRI data. The model yields population bursts that propagate mainly in the feedback direction. In the following, we poise the model in the metastable state and refer to it as the original model.

**Table 1:**
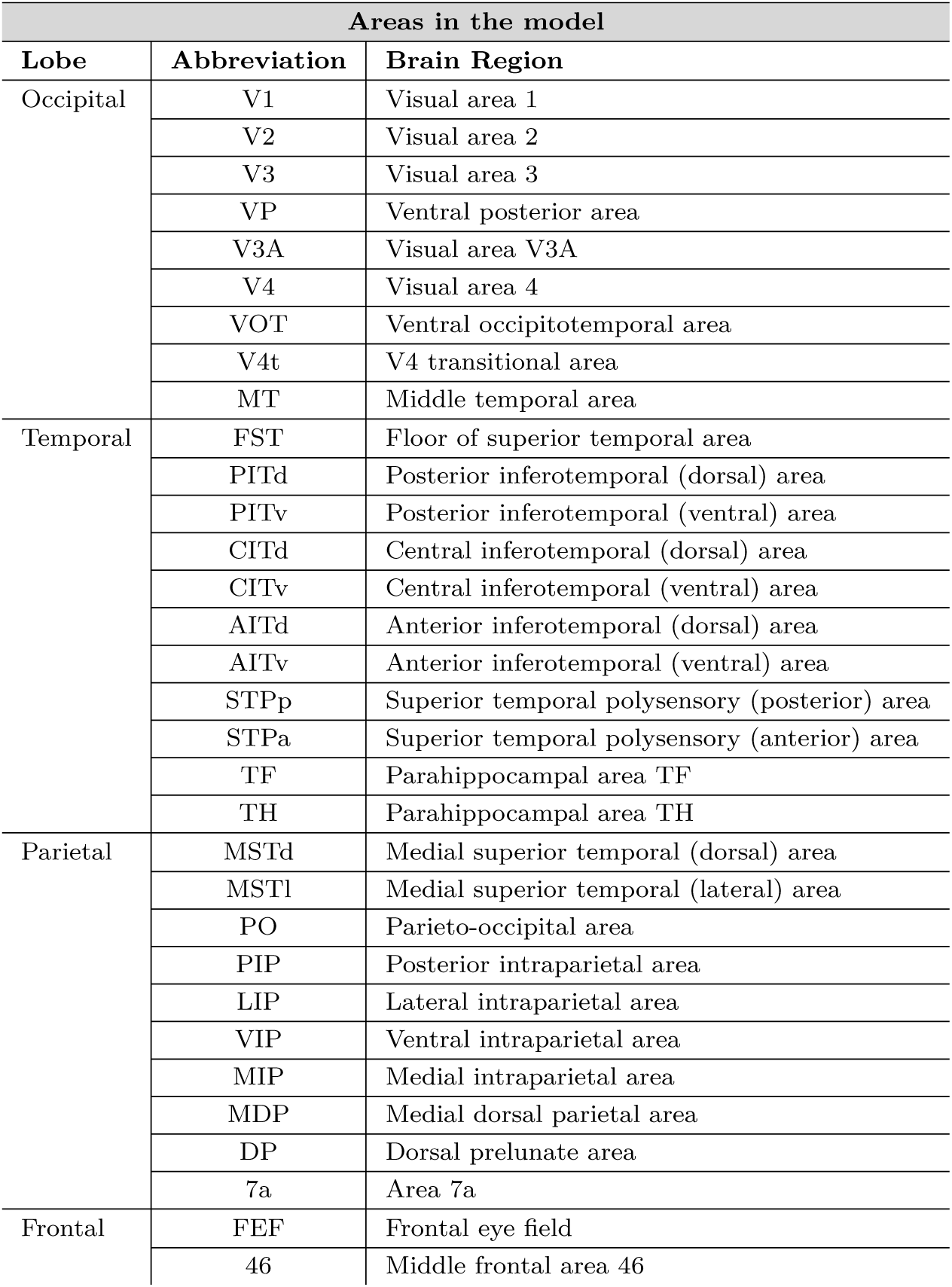
The areas of the model, which include all vision-related areas of macaque cortex in the parcellation of Felleman and Van Essen (1991).

**Table 2:** Model description after Nordlie et al. (2009).

| Model summary |  |
| --- | --- |
| Populations | Original model: 254 populations: 32 areas (Table 1) with eight populations each (area TH: six)<br>Clustered model: Each population is further subdivided into $Q$ clusters. |
| Topology | — |
| Connectivity | area- and population-specific but otherwise random |
| Neuron model | leaky integrate-and-fire (LIF), fixed absolute refractory period (voltage clamp) |
| Synapse model | exponential postsynaptic currents |
| Plasticity | — |
| Input | independent homogeneous Poisson spike trains |
| Measurements | spiking activity |
| Populations |  |
| Type | Cortex |
| Elements | LIF neurons |
| Number of populations | 32 areas with eight populations each (area TH: six), two per layer |
| Population size | $N$ (area- and population-specific) |
| Connectivity |  |
| Type | source and target neurons drawn randomly with replacement (allowing autapses and multapses) with area- and population-specific connection probabilities |
| Weights | fixed, drawn from normal distribution with mean $J$ such that postsynaptic potentials have a mean amplitude of 0.15 mV and standard deviation $\delta J = 0.1J$; 4E to 2/3E increased by factor 2 (cf. Potjans and Diesmann, 2014); weights of inhibitory connections increased by factor $g$; excitatory weights $< 0$ and inhibitory weights $> 0$ are redrawn; cortico-cortical weights onto excitatory and inhibitory populations increased by factor $\chi$ and $\chi_I\chi$, respectively |
| Delays | fixed, drawn from Gaussian distribution with mean $d$ and standard deviation $\delta d = 0.5d$; delays of inhibitory connections factor 2 smaller; delays rounded to the nearest multiple of the simulation step size $h = 0.1$ ms, inter-area delays drawn from a Gaussian distribution with mean $d = s/v_t$, with distance $s$ and transmission speed $v_t = 3.5$ m/s (Girard et al., 2001); and standard deviation $\delta d = d/2$, distances determined as the median of the distances between all vertex pairs of the two areas in their surface representation in F99 space, a standard macaque cortical surface included with Caret (Van Essen et al., 2001), where the vertex-to-vertex distance is the length of the shortest possible path without crossing the cortical surface (Bojak et al., 2011) (see Schmidt et al. (2018a)), delays $< 0.1$ ms before rounding are redrawn |
| Neuron and synapse model |  |
| Name | LIF neuron |
| Type | leaky integrate-and-fire, exponential synaptic current inputs |
| Subthreshold dynamics | $\frac{dV}{dt} = -\frac{V-E_L}{\tau_m} + \frac{I_s(t)}{C_m}$, if $(t > t^* + \tau_r)$<br>$V(t) = V_r$, else<br>$I_s(t) = \sum_{i,k} J_k e^{-(t-t_i^k)/\tau_s} \Theta(t - t_i^k)$, $k$: neuron index, $i$: spike index |
| Spiking | If $V(t-) < \theta \wedge V(t+) \geq \theta$<br>1. set $t^* = t$, 2. emit spike with time stamp $t^*$ |
| Input |  |
| Type | Background |
| Target | LIF neurons |
| Description | independent Poisson spikes (for each neuron, fixed rate $\nu_{\text{ext}} = \nu_{\text{bg}} k_{\text{ext}}$ with average external spike rate $\nu_{\text{bg}} = 10$ spikes/s and number of external inputs per population $k_{\text{ext}}$, weight $J$) |
| Measurements |  |
| Spiking activity |  |

**Table 3:**
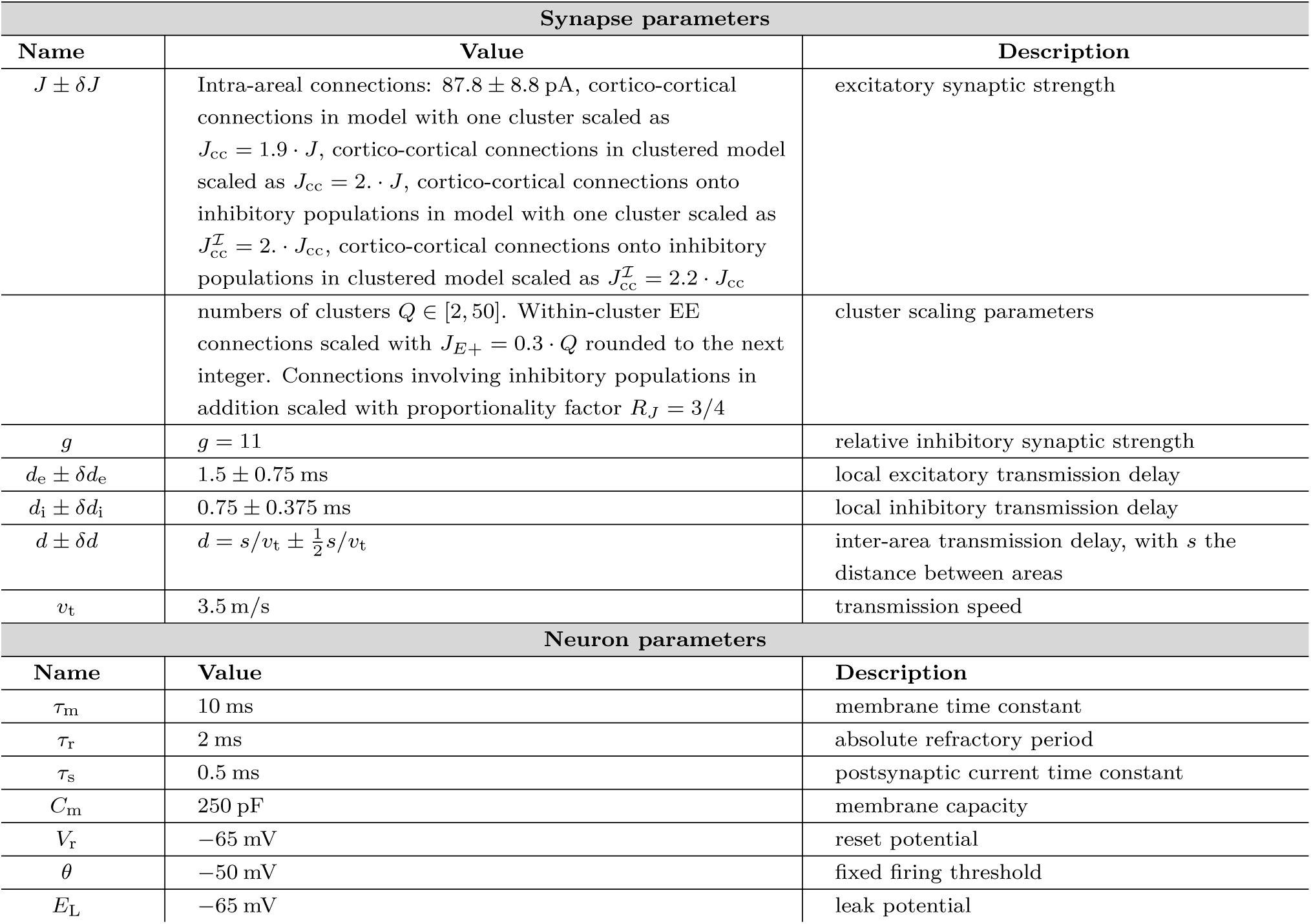
Parameter specification for synapses and neurons.

We extend this model by providing the possibility of injecting a stimulus of variable length and strength into any area. The stimulation consists of spike trains drawn from Poisson processes. The multi-area model distinguishes between input stemming from modeled neurons inside and across areas, and input originating outside the simulated circuitry—that is, the rest of cortex and subcortical structures. The latter input is the background activity driving the multi-area model. We assume that this input becomes stronger during stimulation and thus use the corresponding connections to stimulate the model. The spike trains representing the stimulation are drawn from Poisson processes with stationary rate *ν_stim_* = 30 spikes*/*s and are independent to each target neuron.

### Clustered multi-area model of macaque visual cortex

We generalize a connectivity scheme previously studied in binary (Rost et al., 2018) and spiking networks (Rostami et al., 2022) of one excitatory and one inhibitory population. Our basic building blocks are the layer-resolved microcircuits representing each area. We subdivide these basic building blocks into *Q* equally sized clusters spanning all layers of cortex. Each cluster thus consists of four excitatory and four inhibitory populations, an excitatory-inhibitory pair of populations for each layer. Within a cluster, the excitatory-to-excitatory (EE) synaptic connections are potentiated by a factor *J_E_*_+_. The excitatory-to-inhibitory (EI), inhibitory-to-excitatory (IE), and inhibitory-to-inhibitory (II) synaptic connections are potentiated by a factor *J_I_*_+_. Across clusters, the EE connections are depressed by a factor *J_E−_*, whereas the EI, IE, and II connections are depressed by a factor *J_I−_*. Additionally, a proportionality factor *R_J_*= 3*/*4 is introduced to help prevent firing rate saturation in up states. The factors are related as follows:

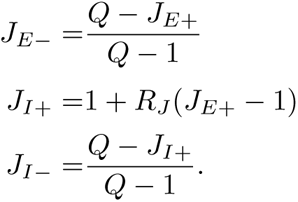

Furthermore, every local cluster has a matching cluster in all other areas that it connects to. The weights between these clusters are scaled as within-cluster weights. Conversely, all cortico-cortical weights between non-matching clusters are scaled as across-cluster weights. A sketch of the network is given in Fig. 1. Following Schmidt et al. 2018b, just as in the original model, we scale cortico-cortical weights onto excitatory populations with a factor *χ* and cortico-cortical weights onto inhibitory populations with a factor *χ_I_χ*. In the case of one cluster, which corresponds to the network studied in Schmidt et al. (2018b), the scaling parameters are *χ* = 1.9 and *χ_I_* = 2. In all simulations involving clusters, we use *χ* = 2. and *χ_I_* = 2.2. The parameters are given in Table 3. Just as in the original model, we draw independent spike trains from Poisson processes with a stationary rate *ν_stim_* = 30 spikes*/*s representing subcortical input to stimulate the model for variable length and strength. However, we selectively stimulate one cluster instead of the whole area. Unless stated otherwise, we report results obtained for the model with *Q* = 50 clusters and refer to it as the clustered model.

### Network simulations

We use commit *c*690*b*7*a* of the NEST 3.0 release (Hahne et al., 2021) running on the JURECA-DC cluster (Thörnig and von St. Vieth, 2021), which hosts compute nodes consisting of two sockets. Each socket contains a 64-core AMD EPYC Rome 7742cd processor clocked at 2.2 GHz equipped with 512 GB of DDR4 RAM. An InfiniBand HDR100/HDR network connects the compute nodes. The simulations are performed using 6 compute nodes with 8 MPI processes each and 16 threads per MPI process. With this setup, building the original model takes 1 minute, whereas the clustered model takes 7 hours. A second of biological time of the original model can be simulated in 165 s. In contrast, the clustered model takes 4 minutes per biological second. In all simulations, time steps of 0.1 ms are used, and the subthreshold dynamics of the leaky integrate-and-fire neuron model is exactly integrated (Plesser and Diesmann, 2009). All presented simulations were run for 101.5 s, of which the first 500 ms are disregarded. In all simulations, spike times were recorded.

### Experimental data

#### Spiking data from macaque cortical areas V1 and V4 in layers 5/6

Neuronal activity was recorded from visual areas V1 and V4 (*N* = 1 subject, *N* = 1 session of *∼* 20 min). Chronic recordings were made using 16 Utah arrays with 8 *×* 8 electrodes each (Blackrock microsystems), 2 of them in visual area V4 and the rest in V1, with a total of 1024 electrodes. The electrodes were 1.5 mm long and thus reached deep layers 5 and 6. The recordings were made in the resting state. The macaque was head-fixed but free to move its limbs, look around and open or close its eyes. Thus, the spiking statistics include data from a few different behavioral states. A full description of the experimental setup, the data collection and preprocessing has already been published (Chen et al., 2022). The raw data were spike-sorted using a semi-automatic workflow with Spyking Circus—a free, open-source, spike-sorting software written in Python (Yger et al., 2018). Multielectrode recordings are prone to cross-talk in the signals leading to above-chance synchronous spiking events. All single units suspected to be cross-talk artifacts were removed from further analysis (Oberste-Frielinghaus et al., 2024). The spiking data were previously published elsewhere (Morales-Gregorio et al., 2023).

#### Spiking data from macaque cortical areas V4 and FEF in layers 2/3

Neuronal activity was recorded from visual area V4 and dorsolateral prefrontal cortex (dlPFC), specifically a part of the frontal eye field (*N* = 1 subject, *N* = 59 sessions of ∼5 min). Acute recordings were made with up to four simultaneous Plexon electrodes, recording from the superficial layers (L2/3) during resting state. The macaque was free to move its limbs, look around and open or close its eyes. Thus, the spiking statistics include data from a few different behavioral states. Spike sorting identified 4*−*10 clean single units per area and session. Single units suspected to be cross-talk artifacts were removed from further analysis (Oberste-Frielinghaus et al., 2024). These data correspond to recordings before or after the behavioral task published in (Sapountzis et al., 2022), using the same recording apparatus.

#### Spiking data from macaque cortical areas DP and 7a in layers 5/6

Neuronal activity was recorded from V1, V2, DP, 7a, and motor cortex (*N* = 1 subject, *N* = 2 sessions of *∼* 10 min). The macaque was implanted with five Utah arrays (Blackrock microsystems), one in V1, one in V2, one in dorsal prelunate cortex (area DP), one in area 7a and one in the motor cortex (M1/PMd). In this study, we only included the 6 *×* 6 electrode arrays from DP and 7a since not enough spikes could be detected in V1 and V2. The electrodes were 1.5 mm long and thus recorded from the deep layers (L5/6) of the cortex. The recordings were made in the resting state. The macaque was free to move its limbs, look around and open or close its eyes. Thus, the spiking statistics include data from a few different behavioral states. The raw signals were spike sorted using the Plexon software. Single units suspected to be cross-talk artifacts were removed from further analysis (Oberste-Frielinghaus et al., 2024). The recording apparatus is described elsewhere (de Haan et al., 2018).

#### Datasets used in Churchland et al. (2010)

Churchland et al. (2010) analyze the Fano factor in seven cortical areas of the macaque monkey: V1, V4, MT, LIP, PRR, PMd, and OFC. In the following, we only consider the first four, as these are part of the multi-area model. Area MT was studied in four different experiments, of which two involved two stimulations. We focus on the experiments in which only one stimulus was applied. The V1 data was taken from an anesthetized monkey, which was presented a 100% contrast sine-wave grating drifting in one of twelve directions. V4 data is taken from a task where the stimulation consisted of one or two oriented bars placed in the neuron’s receptive field. In some experiments, similar bars were placed in the opposite hemifield. The first MT task involved square-wave gratings superimposed to produce a plaid as a visual stimulus. The second stimulation consisted of 0% coherence random dots. The LIP stimulation consisted of two colored saccade targets from which the monkey could choose. To compare the experimental with the simulated data, we extract the experimental Fano factors from Figure 3 in Churchland et al. (2010) using the tool WebPlotDigitizer^1^.

### Analysis methods

#### Summary statistics of the resting-state spiking activity

We use several standard metrics to characterize the resting-state neural activity and to compare the models and experimental data. We use the coefficient of variation of the inter-spike interval distribution (CV ISI) and revised local variation (LvR; Shinomoto et al., 2009) to characterize interval statistics. The CV ISI is defined as the ratio of the standard deviation *σ* to the mean *µ* of the inter-spike intervals,

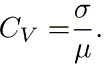

The LvR is defined as

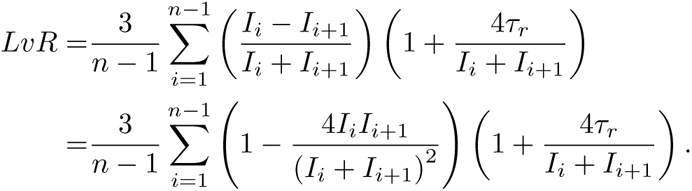

The first term computes the local variance of consecutive inter-spike intervals *I_i_*while the second term accounts for the refractoriness of the neuron.

The distribution of spike rates *P* (rate) is calculated as the histogram of spike rates of all spike trains, computed as the total number of spikes of each neuron divided by the simulation time.

For the experimental data, the CV, LvR, and *P* (rate) are calculated for all spike trains. Where the simulated data are compared against experimental data, as many spike trains as there are in the experimental data are used from randomly drawn neurons. The metrics are then calculated for this subset. This procedure is repeated 100 times, and the mean and standard deviation are calculated. Otherwise, all spike trains from the population are used.

#### Response latency measurement via Poisson surprise

We derive response latencies in a set of areas resulting from an input to V1 employing the Poisson surprise method. This method was first used by Legendy and Salcman (1985) and further refined by Hanes et al. (1995) and Thompson et al. (1996). For a given neuron with mean firing rate *r*, this method evaluates how improbable it is that a series of *n* spikes, which we call response, in a given time interval *T* occurs by chance. The probability *P* is calculated using Poisson’s formula

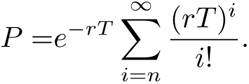

The surprise index

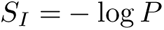

serves as a measure of improbability and yields higher values the more unexpected, or improbable, a result is. In order to detect the response latencies in a given area, we apply the following algorithm to every neuron in the area and identify neurons that exhibit a clear response in at least 60% of the trials. This procedure finds the neurons that reliably respond to the provided stimulus and corresponds to experimentalists probing for responsive neurons. We follow the procedure described by Hanes et al. (1995):

1. We calculate the mean firing rate *r* the neuron for the whole simulation time.
2. We split the spike train into trials, spanning the time between each stimulus onset. For each trial, starting with the first spike, we search for the first two consecutive spikes with a mean firing rate *r*∼ greater or equal to *r*. Then, the first two spikes remain fixed, the next spike is added to the sequence of spikes and the surprise index *S_I_* is calculated. This is done until reaching the end of the trial. The spike where *S_I_*is maximized is defined as the end of the assumed response.
3. To detect responses, we first follow Legendy and Salcman (1985), fixing the last spike of the assumed response and calculating the surprise index *S_I_* for all previous spikes. The spike where the surprise index *S_I_* is maximized is defined as the first spike of the assumed response. Some cortical areas have more gradual responses than the bursting responses considered by Legendy and Salcman (1985). To account for this, Hanes et al. (1995) extended the method from Legendy and Salcman (1985) to determine when a nonbursting change in the activity of the neuron becomes significantly different from the expected Poisson distribution. We follow Hanes et al. (1995) to detect such slow-rising responses. Spikes are added before the one maximizing the surprise index until the surprise index *S_I_* falls below a reduced significance threshold or the algorithm reaches the first spike of the trial. The significance level is set to *p <* 0.01 and relaxed to *p <* 0.05 in areas 7a, MT, MSTl, TF, and FEF. The relaxation is necessary as responses in these areas otherwise go mostly undetected. The assumed response is rejected if its surprise index *S_I_*is not significant.

#### Neural variability quantification across trials

The Fano factor is a measure of the variability of spike trains. It is defined as the ratio of the variance and the mean of the spike counts and measures the response variability across repetitions of the same experimental task, that is, across trials. While its definition is transparent, its results might suffer from careless use (Churchland et al., 2010; Rajdl et al., 2020): At higher spiking rates, the refractory periods of the neurons tend to regularize spiking, which could lower the Fano factor due to the variability of the spiking noise being reduced. Furthermore, the across-trial firing-rate variability could be constant but become normalized by a higher mean after stimulus onset. To control for the influence of the firing rates, we apply the mean matching procedure described by Churchland et al. (2010).

The mean matching procedure works as follows: First, for each neuron, we compute the mean and the variance of the spike counts in a sliding window. Second, we construct the greatest common distribution based on the mean and the variance. The bins of this common distribution have a height equal to the smallest value for that bin across all distributions at all times. Third, at each time, individual points consisting of the mean and variance of the spike counts are excluded until matching the common distribution. Fourth, based on the remaining points, we calculate the Fano factor. Like Churchland et al. (2010), we use a 50 ms wide sliding window that moves in steps of 10 ms.

## Acknowledgements

This project received funding from the DFG in RTG 2416 “MultiSenses-MultiScales” and Priority Program 2041 “Computational Connectomics” [AL 2041/1-1]; and the EU’s Horizon 2020 Framework Grant Agreement No. 785907 (Human Brain Project SGA2) and No. 945539 (Human Brain Project SGA3). A subset of the experimental data was made available in the context of the FLAG-ERA grant PrimCorNet. The authors gratefully acknowledge the computing time granted by the JARA Vergabegremium and provided on the JARA Partition part of the supercomputer JURECA at Forschungszentrum Jülich (computation grant JINB33).

## Author contributions

Conceptualization JP, SvA; Data curation JP, AMG; Formal Analysis JP; Investigation JP; Methodology JP, VR, SvA; Software JP; Visualization JP, AMG; Writing – original draft JP; Writing – review & editing JP, AMG, VR, SvA; Supervision SvA; Funding acquisition SvA

## Supplementary materials

### Supplementary methods

#### V1 spiking data from Chu et al. (2014b)

The experimental recordings from all layers of V1 have previously been used and described in the context of the multi-area model in Schmidt et al. (2018b). A detailed description has been published in Chu et al. (2014b), and the dataset is publicly available (Chu et al., 2014a). In short, the data were collected from a 64-electrode array implanted into primary visual area V1 of a lightly anesthetized macaque monkey, and are spike-sorted into 140 single units. In our analysis in Fig. S1, we used the data on 15 minutes of spontaneous activity during which no visual stimulation was provided.

#### Macaque resting-state fMRI

The fMRI data have previously been used and described in the context of the multi-area model in Schmidt et al. (2018b). The data are publicly available in processed form in the GitHub repository of the model^2^. The data were acquired from six male macaque monkeys, and five of the six subjects have previously been described in Babapoor-Farrokhran et al. (2013). The Animal Use Subcommittee of the University of Western Ontario Council on Animal Care approved all experimental protocols in accordance with the guidelines of the Canadian Council on Animal Care. The subjects were under light anesthesia, and ten sets of five-minute resting-state fMRI scans were acquired from each subject. The AFNI software package^3^ was used to regress out nuisance variables. The Pearson correlation coefficients of the probabilistically weighted ROI time series for each scan were used to compute the functional connectivity (Shen et al., 2012).

#### Power spectral density comparison

For the comparison with the V1 data in Fig. S1, the power spectral density (PSD) is computed using Welch’s method implemented in signal.welch in the Python SciPy library (Virtanen et al., 2020). A boxcar window, a segment length of 1024 data points, and 1000 overlapping points between segments are used.

#### Functional connectivity comparison with fMRI

For the analysis of the functional connectivity (FC) in Fig. S1, we define the FC of the spiking network model as the zero-time-lag cross-correlation of the area-averaged synaptic inputs, following Schmidt et al. (2018b). It is approximated by

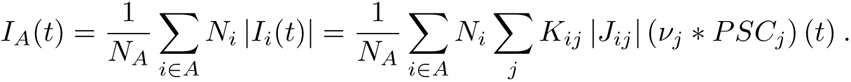

The term *PSC_j_*(*t*) = exp[*−t/τ*_s_] is the normalized postsynaptic current, *∗* means convolution, *τ*_s_ is the synaptic time constant, *ν_j_* is the population firing rate of the source population *j*, *K_ij_* is the mean indegree, and *J_ij_* is the mean synaptic weight of the connection from *j* to the target population *i* containing *N* neurons. The population firing rate *ν_j_* is a spike histogram with bin width 1 ms averaged over the entire population. Hence, time *t* here has a resolution of 1 ms.

#### Supplementary results

##### Comparison with Schmidt et al. (2018b)

To understand the main differences between the original and clustered model, we first compare the simulation results against the experimental data used by Schmidt et al. (2018b). The experimental data consist of multielectrode spike recordings from macaque V1 (Chu et al., 2014b,a) and resting-state fMRI recordings (Babapoor-Farrokhran et al., 2013); see Experimental data for a detailed description of the data. Fig. S1 shows that the explanatory power of the model is conserved. Fig. S1A–C show raster plots of the original model, the clustered model, and the experimental recordings of (Chu et al., 2014a). The CV ISI, shown in Fig. S1D, and the LvR, shown in Fig. S1E, in area V1 appear slightly better in the clustered model than in the original model.

We quantify the similarity between the distributions using a Kolmogorov-Smirnov test of each model against the experimental data distribution. The CV ISI distribution of the V1 activity in the clustered model appears more similar to the experimental data (KS = 0.53, p *«* 0.001) than the original model (KS = 0.73, p *«* 0.001).

Likewise, the LvR distribution is also better captured by the clustered model (KS = 0.59, p *«* 0.001) than by the original model (KS = 0.64, p *«* 0.001). However, the firing rate distribution from the original model better matches the experimental data (KS = 0.09, p = 0.28) than the clustered model (KS = 0.26, p *«* 0.001), shown in Fig. S1F. We also compare the power spectral density (PSD) of the spike histograms (bin size of 1 ms). The original model matches the experimental power spectrum well, whereas the clustered model exhibits an almost flat power spectrum that does not follow the experimental findings.

Finally, we compare the cortico-cortical interactions of the model with experimentally measured fMRI BOLD signals (Babapoor-Farrokhran et al., 2013). Fig. S1H shows the Pearson correlation coefficient *r* of simulated functional connectivity (FC) with experimentally measured FC for different numbers of clusters *Q*. The dashed line shows the average correlation coefficient (*r* = 0.31) across all monkeys in the fMRI dataset, which indicates the correlation level to be explained by a model that is not tuned to any individual subject. All model versions reach this level of correlation between simulated and experimental FC.

**Figure S1:**
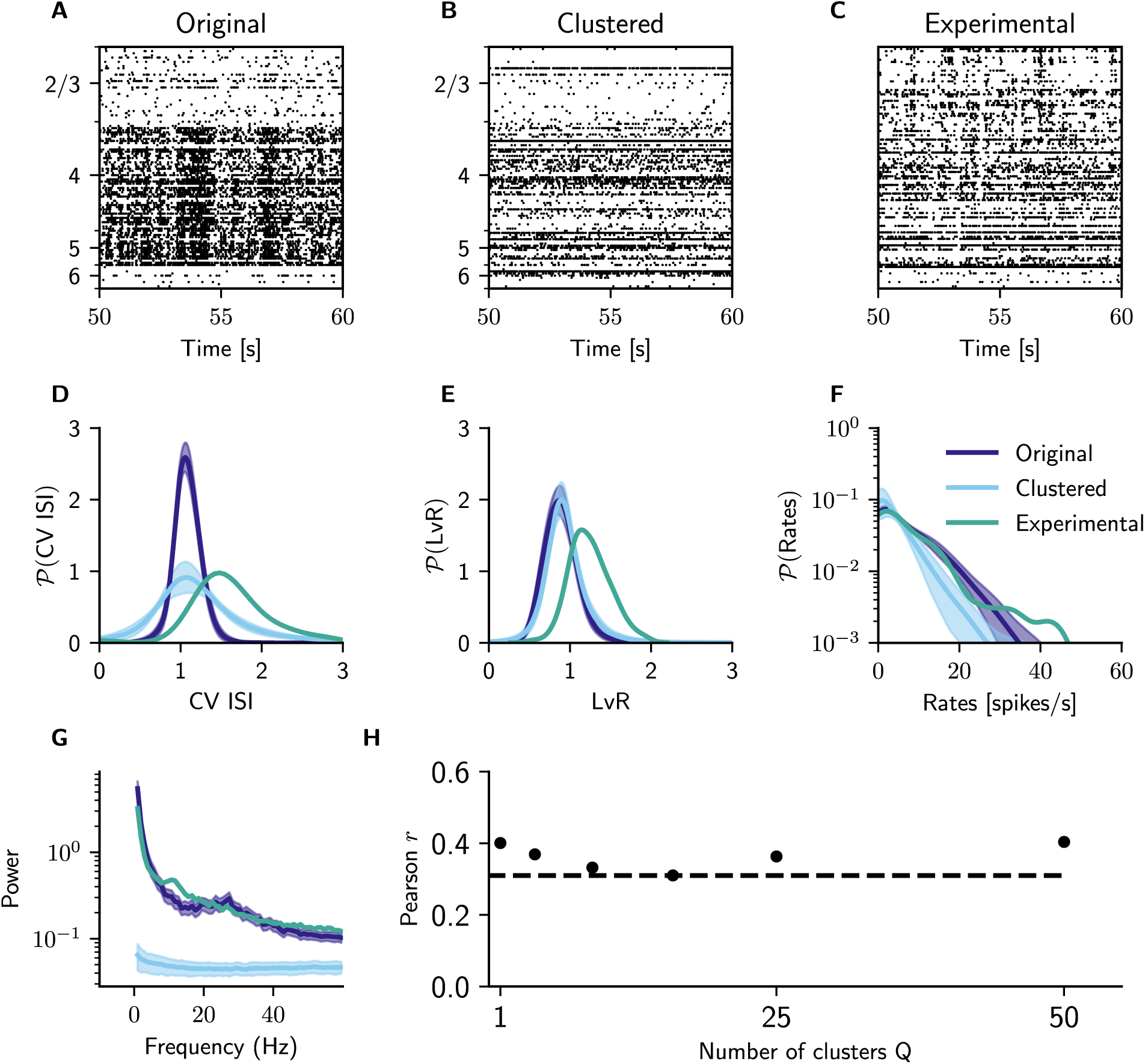
Comparison with experimental data used in Schmidt et al. (2018b). (A-C) Raster plots showing the spiking activity of excitatory and inhibitory cells from the original, unclustered model **(A)**, from the clustered model with 50 clusters **(B),** and from the experimental data **(C);** simulated excitatory and inhibitory neurons are shuffled within layers for plotting to better match the sampling from the experimental data, where neurons are ordered depth-wise but no distinction is made between excitatory and inhibitory cells. **(D,E)** Distribution of irregularity of single-unit spike trains across all populations in area V1 quantified by the coefficient of variation of the interspike intervals *CV ISI* **(D)** and revised local variation *LvR* **(E)** (Shinomoto et al., 2009) for different numbers of clusters *Q* compared against experimental data (Chu et al., 2014a). **(F)** Distribution of simulated spike rates across all populations of V1 and for the 140 single units extracted from the experimental data (Chu et al., 2014a). **(G)** Power spectra of the summed spiking activity of 140 randomly selected V1 neurons for the two model versions and for the experimental data. **(H)** Pearson correlation coefficient *r* of simulated functional connectivity FC vs. experimentally measured FC for different numbers of clusters. *Q* = 1 refers to the original, unclustered model. Dashed line: average correlation across all monkeys.

## Footnotes

1 https://automeris.io/WebPlotDigitizer/

2 https://github.com/inm-6/multi-area-model

3 fni.nimh.nih.gov/afni

